# A survey of ADP-ribosyltransferase families in the pathogenic *Legionella*

**DOI:** 10.1101/2024.07.30.605764

**Authors:** Marianna Krysińska, Marcin Gradowski, Bartosz Baranowski, Krzysztof Pawłowski, Małgorzata Dudkiewicz

## Abstract

**Background:** ADP-ribosyltransferases (ARTs) are a superfamily of enzymes implicated in various cellular processes, including pathogenic mechanisms. The *Legionella* genus, known for causing Legionnaires’ disease, possesses diverse ART-like effectors. This study explores the proteomes of 41 *Legionella* species to bioinformatically identify and characterise novel ART-like families, providing insights into their potential roles in pathogenesis and host interactions.

**Methods:** We conducted a comprehensive bioinformatic survey of 41 *Legionella* species to identify proteins with significant sequence or structural similarity to known ARTs. Sensitive sequence searches were performed to detect candidate ART-like families. Subsequent validation, including structure prediction of such families, was achieved using artificial intelligence-driven tools, such as AlphaFold. Comparative analyses were performed to assess sequence and structural similarities between the novel ART-like families and known ARTs.

**Results:** Our analysis identified 63 proteins with convincing similarity to ARTs, organised into 39 ART-like families, including 26 novel families. Key findings include:

- DUF2971 family: exhibits sequence similarity to DarT toxins and other DNA-acting ARTs.
- DUF4291 family: the largest newly identified family shows structural and sequence similarity to the diphtheria toxin, suggesting the ability to modify proteins.

Most members of the novel ART families are predicted effectors. Although experimental validation of the predicted ART effector functions is necessary, the novel ART-like families identified present promising targets for understanding *Legionella* pathogenicity and developing therapeutic strategies. We publish a complete catalogue of our results in the astARTe database: http://bioinfo.sggw.edu.pl/astarte/.

## Introduction

ADP-ribosyltransferases (ARTs) are enzymes that chemically modify proteins, nucleic acids and small molecules by covalently attaching an ADP-ribose derived from oxidised nicotinamide adenine dinucleotide (NAD⁺) via N-, O- or S-glycosidic bonds (Corda, 2003; Aravind et al., 2015; Cohen & Chang, 2018). ADP-ribosyltransferases are typically classified as monoARTs or poly(ADP-ribose) polymerases (PARPs), catalysing the addition of a single ADP-ribose moiety or polymers of ADP-ribose to its targets. Further, the ADP-ribose chain catalysed by PARPs can be either linear or branched (Ueda & Hayaishi, 1985; Sugimura & Miwa, 1994; Aravind et al., 2015; Munnur & Ahel, 2017).

ADP-ribosyltransferases are common in nature and are found in all domains of life and in viruses. The characteristic feature of the ART superfamily is a conserved structural core formed by a split beta-sheet formed by six or seven antiparallel beta strands and surrounded by alpha helices. As a result, the NAD⁺ molecule is sandwiched between the two halves of the split beta sheet (Aravind & de Souza, 2012; Aravind et al., 2015; Cohen & Chang, 2018). Almost all ADP-ribosyltransferases have additional protein domains accompanying the ART catalytic domain that target enzymes to their substrates and specific cell locations (Vyas et al., 2013; Bock & Chang, 2016; Cohen & Chang, 2018). In contrast to the structural similarity, sequence conservation between ART families is poor, which is also manifested by the diverse amino acid composition of the active sites. Classifying by active site types, the ART superfamily contains four main clades: HHh, RSE, HYE, and the so-called atypical clade (Aravind et al., 2015).

Mono-ADP-ribosylation is believed to have originally appeared in bacteria as a defence mechanism against viruses or bacteria (Aravind et al., 2015; Koch-Nolte, 2015). Poly-ADP-ribosylation emerged in Eukaryotes as a process involved in DNA-repair, modulation of chromatin structure and programmed cell death (Morales et al., 2014).

During infection, ADP-ribosylation of host proteins by bacterial effectors can change the apoptotic potential of the cell, disrupt the organisation of the actin cytoskeleton, alter the organisation of cell membranes, and interfere with the cellular immune response (Maresso et al., 2007; Hottiger et al., 2010; Aravind et al., 2015; Komander & Randow, 2017; Klockgether & Tümmler, 2017; Cohen & Chang, 2018). Other toxins ADP-ribosylate nucleotides in single-stranded DNA (ssDNA). This leads to ADP-ribosylation of the origin of chromosomal replication of DNA and hence inhibition of cell growth (Jankevicius et al., 2016). A similar mechanism is used by bacteria for self-defence during phage infection. For example, DarT from *M. tuberculosis* selectively ADP-ribosylates thymidine nucleotides within phage ssDNA which blocks phage DNA replication (Ritter et al., 2003).

*Legionella* is a genus comprising free-living, biofilm-associated, or host-associated bacteria (Taylor, Ross & Bentham, 2009). *Legionella* infection in humans can cause severe atypical pneumonia called legionellosis or a milder version called Pontiac fever (Fraser et al., 1977; McDade et al., 1977; Brenner, Steigerwalt & McDade, 1979; Newton et al., 2010; Mondino et al., 2020; Kanatani et al., 2021). Currently, the genus *Legionella* comprises more than 70 different species of bacteria, about half of which have been found to be pathogenic to humans. Additionally, the majority are considered potential human pathogens (Mercante & Winchell, 2015; Chambers et al., 2021; Kanatani et al., 2021). After entering the host cell, *Legionella* creates a special vacuole in which it can survive and reproduce called the Legionella Containing Vacuole (LCV) (Samrakandi et al., 2002). Effector proteins translocated by *Legionella* into the host cell can profoundly alter the host cell’s behaviour, which greatly facilitates *Legionella* replication and pathogenesis. These alterations include safeguarding the LCV from degradation, manipulating the endomembrane system, dampening the host immune response, and exploiting the hosts resources (Segal, Feldman & Zusman, 2005; Isberg, O’Connor & Heidtman, 2009; Burstein et al., 2016; Gomez-Valero et al., 2019). The most thoroughly-characterized *Legionella* species, *L. pneumophila*, translocates over 330 effectors (Burstein et al., 2016; Wexler et al., 2022). Three recently described *L. pneumophila* effectors with ART functions are lpg0181/Lart1 (Black et al., 2021), lpg0080 (Fu et al., 2022) and the SidE family of all-in-one Ub ligases. The first was identified as an ADP-ribosylating factor for a specific class of NAD+-dependent glutamate dehydrogenase (GDH) enzymes found in fungi and protists, which includes many natural hosts of *Legionella* (Black et al., 2021). The second (lpg0080) was identified as a mono-ADP-ribosyltransferase that localises to mitochondria in host cells and ADP-ribosylates the ADP/ATP translocase, which impairs its activity (Fu et al., 2022). The third (SidE family) shows atypical activity by attaching phosphoribosyl ubiquitin (PR-Ub) to serine residues on substrates, using the phosphodiesterase and mono-ADP-ribosyltransferase domains. SidE proteins modify small Rab GTPases, disrupting vesicle movement and cell membrane dynamics, preventing the maturation of phagosomes into bacterial lysosomes, and thus allowing *Legionella* to establish a replication niche (Akturk et al., 2018).

In this article, basing on our expertise in identifying novel enzyme families (Black et al., 2021; Wyżewski et al., 2021), and motivated by many known bacterial effectors and toxins being ARTs (Black et al., 2021; Fu et al., 2022), we undertook a bioinformatics search and survey of novel ART-like domains in 41 *Legionella* proteomes. We identified 26 novel ART-like families, six of which we describe in detail. Presenting a catalogue of ART-like catalytic domains in *Legionella,* we compare the known and the novel families, predict biological functions for the novel ones, establish their evolutionary history and occurrence across the bacterial world.

**Figure 1.**
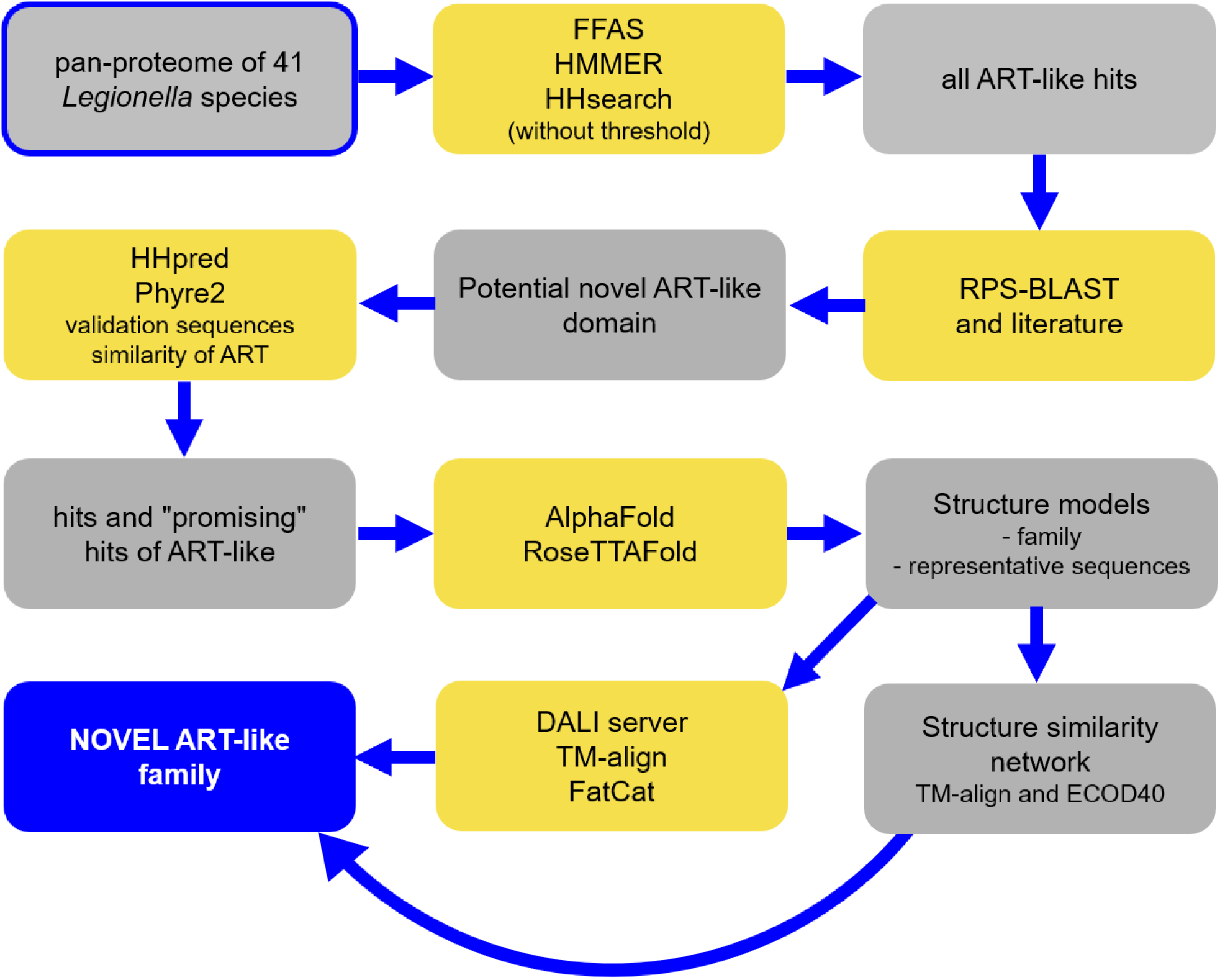
Methods used to identify new ARTs in *Legionella* sp.; grey rectangles – data, yellow rectangles – tools used.

## Materials and methods

### Sequence data

Proteomes for 41 species of Legionella were provided by the study by Burstein et al. (Burstein et al., 2016). To reduce the computational burden, the sequences were clustered by sequence similarity in two steps using the CD-HIT algorithm with sequence identity cut-offs 70% and 50% (Li, Jaroszewski & Godzik, 2001, 2002; Li & Godzik, 2006; Huang et al., 2010). Next, 21,616 resulting protein sequences from 41 *Legionella* proteomes were cut into fragments of 300 amino acids with an overlap of 100 amino acids. This method facilitates the detection of domains in multi-domain proteins, while using an overlap can help detect domains at fragment boundaries.

### Remote homology detection

For distant similarity recognition to ART families, four methods were used: 1) the profile-profile alignment and fold recognition algorithm – FFAS (Xu et al., 2014) (searching the COG, Hsapiens, PDB, SCOP, ECOD and Pfam databases), 2) homology detection and structure prediction method HHpred (Gabler et al., 2020) and HHsearch pipeline (databases searched: Pfam, PDB, SCOP) that uses hidden Markov model HMM-to-HMM comparison; 3) a similar method, Phyre2 (Kelley et al., 2015), which additionally models 3D structure of query and compares it with 3D models library, and 4) homology detection by comparing a profile-HMM to either a single sequence or a database of sequences – HMMER algorithm (Eddy, 2011). Standard parameters and significance thresholds were selected except for the HHpred algorithm, where several parameters have been modified: MSA (Multiple Sequence Alignment) generation method (all options were used); Alignment Mode (local:norealign; local:realign; global:realign); Min probability in hitlist (%) used 10% or 20%; Min coverage of MSA hits (%) used 10% or 20%; MSA generation iteration number was 0, 3, 5 or 8 and E-value cutoff was 1e-3, 0.05 or 0.1 (see Fig. 1 and Suppl. Tables S1-S5).

**Figure 2.**
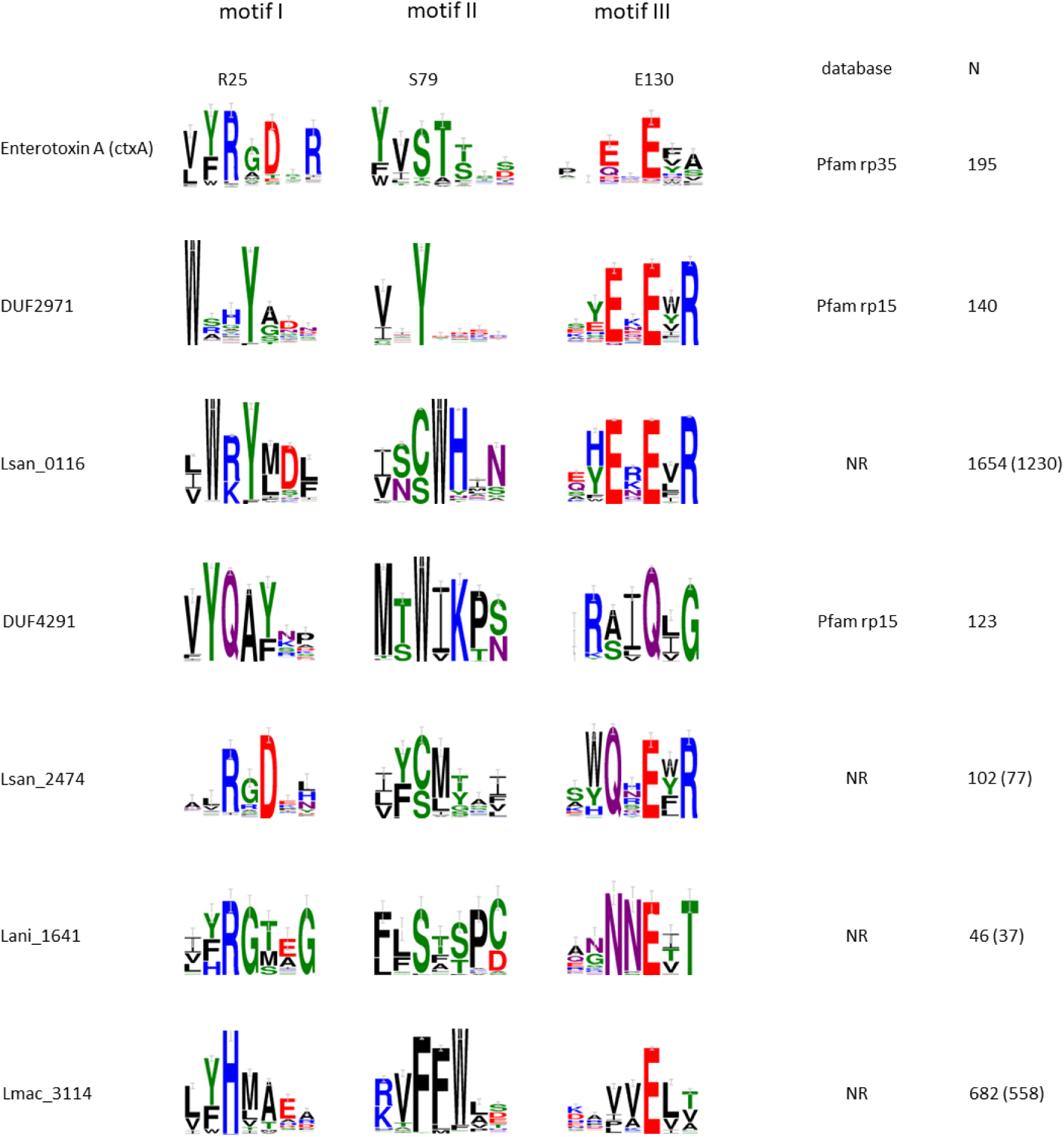
The tripartite active site signature [R/H]x[D/T] - [S/Y][T/x][S/x] - [Q/E]x[Q/E/D] is well conserved in most new ART families. Active site motif sequence logos for selected families. An archetypal ADP-ribosyltransferase, Enterotoxin a, ctxA, is shown at the top. Numbering of the residues (top row) according to ctxA sequence (O’Neal et al., 2004). The DB column indicates the source database of ART sequences used for logos. N is the number of homologous sequences from the BLAST search (E-value = 1e-4). In parentheses - numbers of homologous sequences after CD-HIT clustering at 99% sequence identity.

### Multiple sequence alignments and sequence logos

Members of novel families were manually collected using BLAST (Altschul et al., 1997; Boratyn et al., 2012) and non-redundant sequence database (NR) (Sayers et al., 2021) (E-value = 1e-4 or with default options). In cases, when BLAST found less than 5 homologs, PSI-BLAST was used (3 or 5 iterations with default options) (Boratyn et al., 2012). Multiple sequence alignments were made using the MAFFT algorithm with default settings (Katoh et al., 2002; Katoh, Rozewicki & Yamada, 2019). Next, the sequence logos were prepared using WebLogo3 (Crooks et al., 2004). Colouring according to amino acid chemistry was used. For the logos, the alignments were processed with an in-house script that removes alignment columns that contain gaps in the reference sequence.

### Structure modelling and comparison

The structures of representative proteins from new ART-like families were modelled with the use of two different tools: AlphaFold (Jumper et al., 2021) and RoseTTAFold (Baek et al., 2023). Comparisons of structures were performed with FATCAT (Li et al., 2020), Dali server (Holm, 2020) and TM-align (Zhang & Skolnick, 2005) (Suppl. Table S6 and data made available in the astARTe online database).

### Visual clustering of families (sequence analysis of families relations, Fig. 5)

The CLANS program (Frickey & Lupas, 2004) was used to visualise the relationships between clusters of ADP-ribosyltransferases families. The collection of the known ADP-ribosyltransferase domains was obtained from the Pfam database, as the sets of sequences from families of ADP-ribosyl clan CL0084 – Pfam 35.0 (Mistry et al., 2021) (see Suppl. Table S9B). Then the sequence sets from ART families not yet included in the ADP-ribosyl Pfam clan were added. Next, collected data were supplemented with sequence sets representing the following ART-like domain families not included in the Pfam database (see Suppl. Table S9B) and, in the end, we added all 26 newly predicted ART families with their homologs collected by one of the three methods with cut-off threshold of 99%: 1) BLAST, NR, E-value = 1e-4; 2) PSI-BLAST, two iterations, NR, default; 3) PSI-BLAST, three iterations, NR, default (see Suppl. Table S9C). Additionally, two families DUF2971 (PF11185, rp15 sequence set) and DUF4291, rp15 set were included (see all details in Suppl. Table S9A).

Next, the whole collection of putative and known domain sequences were prepared for the CLANS procedure by clustering each family separately with CD-HIT and selecting representatives at a 70% (known domains) or 99% (putative domains) sequence identity threshold (Huang et al., 2010). CLANS algorithm was run with following parameters: BLOSUM45 (scoring matrix) and E-value = 1 or E-value = 10e-2.

### HMMs of known families for searching sequence databases and profile databases

Sequence sets for HMM construction are collected in a similar manner as the data for CLANS analysis. The most important difference: all sequence sets for families present in the Pfam database were downloaded as rp75 sequence sets from Pfam 34.0 (Mistry et al., 2021) (see all details in Suppl. Table S9B).

In the next step, we rejected short sequences (less than 50 amino acids) and applied a sequence similarity cut-off threshold of 80% (CD-HIT). The ClustalO program (Sievers & Higgins, 2018) was used to build multiple sequence alignments. HMMs were prepared by the hmmbuild program from the HMMER software package (Eddy, 2011).

### Species dendrogram and estimating numbers of homologs

Dendrogram was made from alignment of 16S rRNA sequences for the species in the family *Legionellaceae*. The sequences were downloaded from NCBI (“Nucleotide [Internet]. Bethesda (MD): National Library of Medicine (US), National Center for Biotechnology Information”) and SILVA database (Glöckner et al., 2017). The alignment was performed via the ngphylogeny.fr server (Dereeper et al., 2008; Lemoine et al., 2019) using the Muscle 3.8.31 algorithm with default parameters (Edgar, 2004). A dendrogram was constructed using the PhyML method with the Approximate Likelihood-Ratio Test (aLRT): SH-like (Guindon et al., 2010). Homologs for all families were collected using the hmmsearch algorithm (E-value = 1e-4) (Eddy, 2011) by using the HMMs of known and new ART and ART-like families as queries against the RefSeq database (Sayers et al., 2021) (see all details in Suppl. Table S9C).

Species dendrogram was visualised using the iTol server (Letunic & Bork, 2021).

### Structure similarity network

Sequences for each of the new families were collected (see Suppl. Table S9D) and protein domains were modelled using ColabFold (Mirdita et al., 2022) (an implementation of AlphaFold (Jumper et al., 2021) using the fast homology search of MMseqs2 (Steinegger & Söding, 2017; Mirdita, Steinegger & Söding, 2019)) or ESMFold (Lin et al., 2023). The resulting 432 models and 59 ART structures from the ECOD40 (version 2022/09/12, develop286) database were compared (all to all) using the TM-align algorithm (Zhang & Skolnick, 2005). Based on the obtained TM-score values (nTM-score, score normalised against a smaller structure; with threshold better than 0.3, and processed: *ln(nTM-score)×(−1)*) for each pair of structures (so that the new score is in the range 0-1), a structure similarity network was constructed in CLANS algorithm (see Suppl. Table S9D). We checked the quality of the models by analysing the pLDDT and pTM parameters for each model (see Suppl. Table S9E). The same set of sequences was used for the sequence similarity network (Fig. 4).

### NAD docking to models

We made models (ColabFold) for reference sequences from the 6 new families discussed in this article. NAD docking was performed using the HADDOCK 2.4 (High Ambiguity Driven protein-protein DOCKing) web service (van Zundert et al., 2016; Honorato et al., 2021) package with automatic (CPORT (Vries & Bonvin, 2011)) and manual indication of the binding site residues. Visualisation of the results was done in UCSFChimera (Pettersen et al., 2004) using ConSurf server (Ashkenazy et al., 2010) for mapping MSA conservation into reference sequences models for 6 families. Structures superposition and the closest structural homologues for models were obtained using the DALI server. Additionally, proteins representative of the families described were modelled with AlphaFold3 (Abramson et al., 2024) including NAD as a ligand. The high-quality models obtained (pLDDT>90, pTM>90) confirmed the docking results – location and position of the NAD (data not shown).

### Additional analyses

The subcellular localization was predicted by SubCons (Salvatore, Shu & Elofsson, 2018). Prediction of the presence of signal peptides was made in SignalP 6.0 (Community, 2021). Predictions of effector function were done using EffectiveDB (Arnold et al., 2009; Eichinger et al., 2016) and BastionX (Wang et al., 2021) with standard parameters. Analysis of genomic neighbourhoods was done using the ProFaNa tool (Baranowski & Pawłowski, 2023) and we used the BioCyc portal for detailed analysis of the operons ((Karp et al., 2017)). The occurrence of additional domains was analysed using Batch CD-Search (RPS-BLAST) (Marchler-Bauer et al., 2011).

### Alternative approach to the detection of distantly related homologues

Alternatively, by using RoseTTAFold structure models for Pfam families, we found an ART-like family in *Legionella* that was not detectable by sequence similarity. Here, the pipeline included pairwise TM-align structure alignments between RoseTTAFold models provided in the Pfam 35.0 database and a set of reference ART domain structures from the ECOD database (ECOD40).

### Definition of new families

To define what is meant by “novel family” we applied several criteria. A novel family is a set of proteins not described in the literature as ADP-ribosyltransferases and not annotated such, with similarity to known ARTs not detectable using standard sequence analysis methods – RPS-BLAST or Pfam HMM. Additionally, a family designated as “novel”, based on CLANS clustering analysis, must be sufficiently different from known and other novel ART families.

## Results

### Search for ART superfamily members in *Legionella*

We started our survey with the pan-proteome from 41 *Legionella* species (see Methods). After data pre-processing (see Methods), 21,616 sequences were analysed by FFAS03, HMMER, and HHsearch for sequence similarity to known protein domains. The raw results were filtered by a Python text-mining script that retrieved all ART-like hits using Pfam, PDB, and SCOP identifiers as well as keywords in protein names. In this way, 63 potential ART-like domains were identified exhibiting any similarity to known ART-like domains, including statistically non-significant similarity (Suppl. Tables S2-S4). In the next step, we applied RPS-BLAST validation and literature searches, and known ADP-ribosyltransferases were filtered out. The known hits included 21 proteins from 11 families: RES (Pfam identifier PF08808), PARP (PF00644), ART (PF01129), Dot_icm_IcmQ (PF09475), DarT (PF14487), DUF952 (PF06108), lpg0080, Lart1 and the poorly studied AbiGi family (PF10899), presumably part of the type IV toxin-antitoxin system, as well as the FRG family identified recently by Aravind (Aravind et al., 2015; Burroughs & Aravind, 2020) as an ART-like family present in *Legionella* (Suppl. Tables S1-S2). Forty-two potential ART-like FFAS-HHsearch-HHMER hits were not automatically identified as ART-like and were verified by other distant sequence similarity search methods (Phyre2, HHpred and CLANS analysis) and de novo structure modelling using the RoseTTAFold and AlphaFold methods supplemented with structural comparisons (FATCAT and Dali servers) (Suppl. Tables S5-S6). After discarding 3 false positive ARTs, remaining 39 atypical ART proteins were grouped in twenty-six novel families (see Suppl. Tables S6, S7 and S11).

We identified unequivocal active site signatures in 15 families. In 7 families, we found the presence of conservative substitutions in the predicted active sites (see Fig. 2 and Tab. 1). In the remaining 4 families, the active site is partly non-conserved which suggests they are pseudoenzymes, i.e. proteins that are homologous to active enzymes but presumably lack catalytic activity because of mutations to critical active site amino acids. We classified six of the new families into the HYE clade, one into the HHh clade, one into the atypical clade, and the rest (18 families) into the RSE clade (Fig. 3).

**Figure 3.**
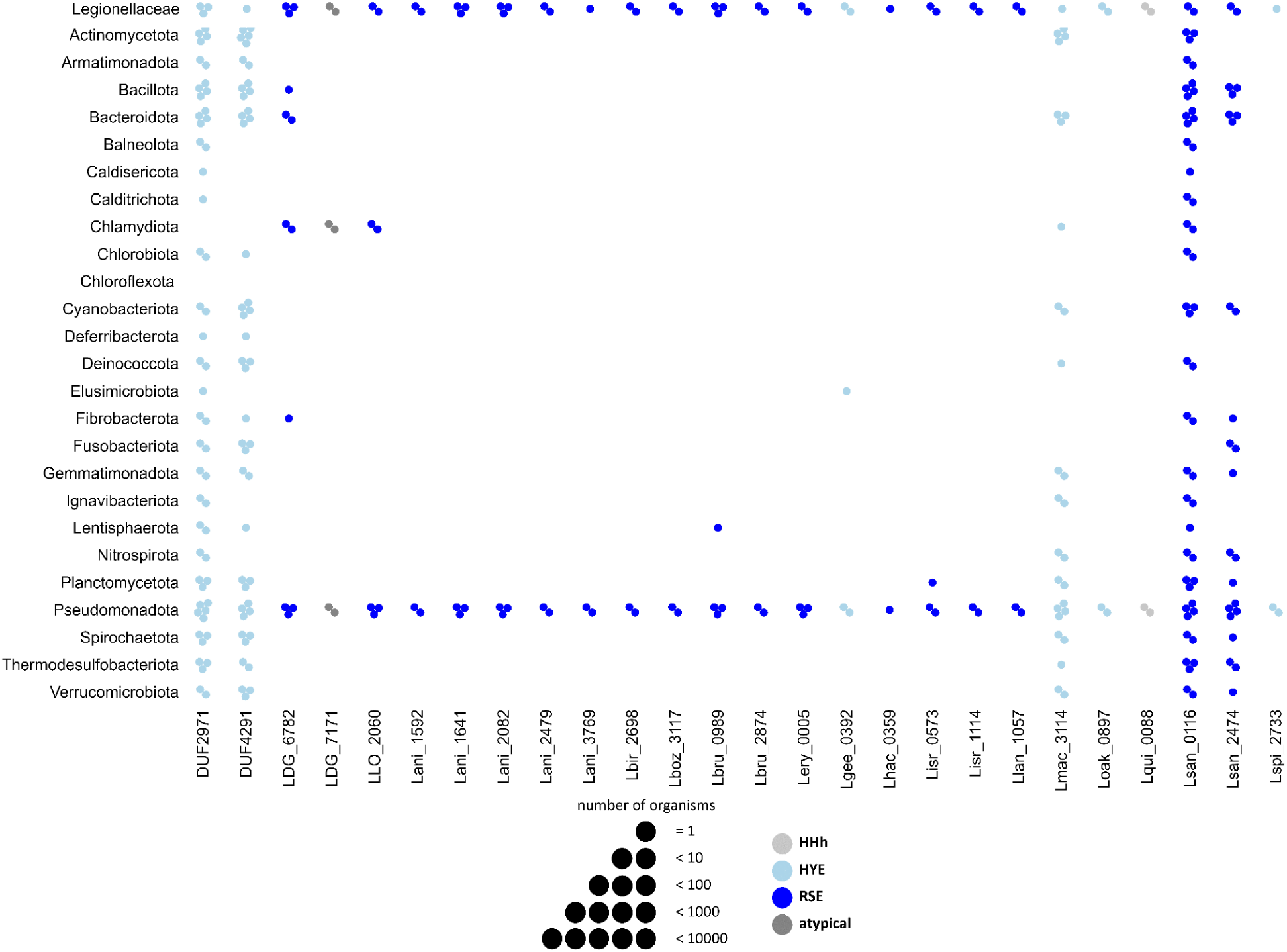
Taxonomic distribution of members of novel ART families in bacterial phyla. The family *Legionellaceae* was considered separately. The number of bacterial strains in which homologues of new families were found is shown on a logarithmic scale (see legend of the figure). Colours reflect ART clade membership.

**Figure 4.**
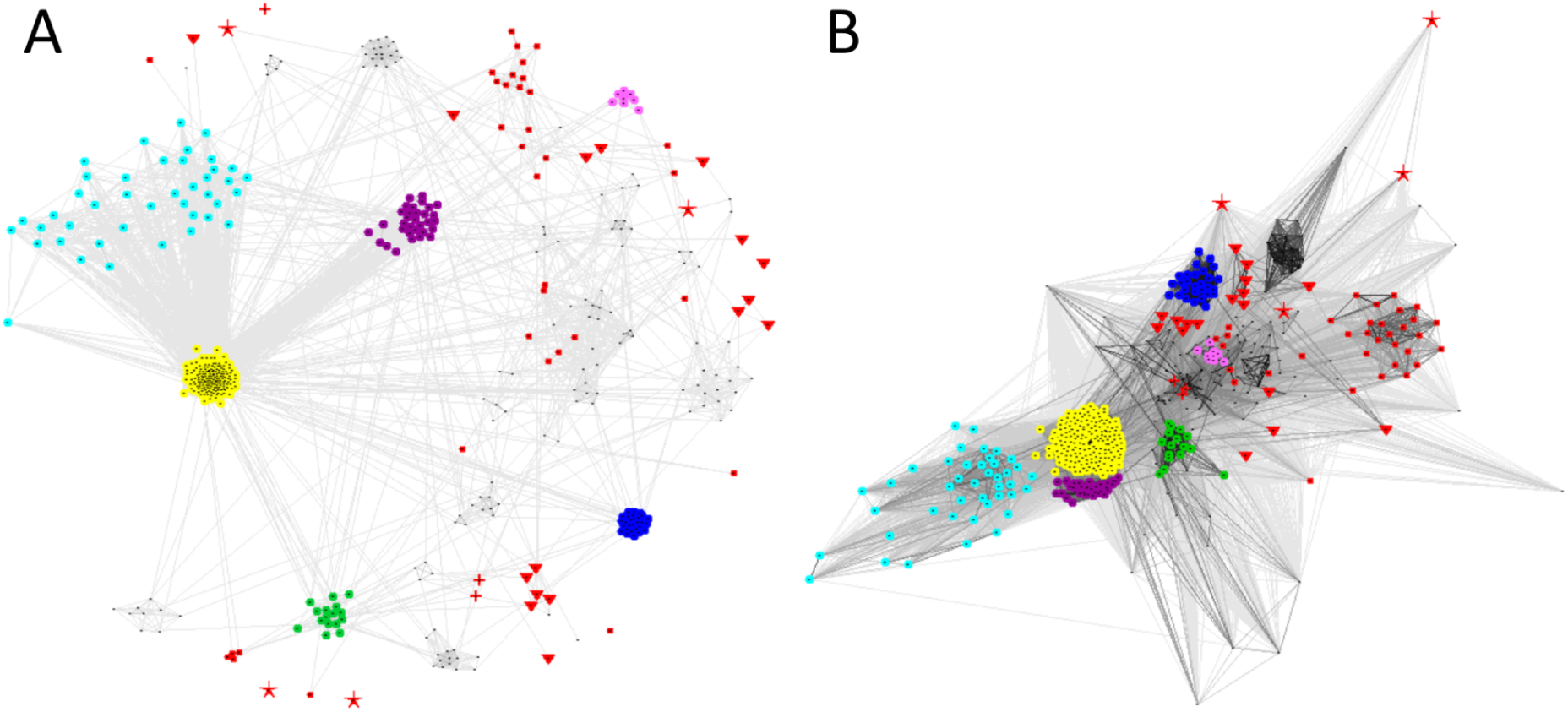
Sequence and structure similarity relationships between novel and known ART families. The CLANS graph shows: (A) the sequence similarities obtained using pairwise BLAST comparisons, taking into account significant and borderline significance similarities up to the E-value of 1 (BLOSUM45); (B) the structure similarities obtained using TM-align all-to-all comparisons (with TMscore threshold better than 0.3). Novel families discussed in detail shown: DUF2971 – turquoise, Lsan_0116 – yellow, DUF4291 – blue, Lsan_2474 – purple, Lani_1641 – pink and Lmac_3114 – green. Known ART families (as defined in the ECOD40 dataset) are marked as red (crosses – clade HHh, dots – clade RSE, triangles – clade HYE, asterisks – atypical clade).

CLANS sequence and structure similarity graphs capture distant relationships between the novel *Legionella* ART-like families and fifty-nine representative known ART domains from ECOD40 database (Cheng et al., 2014) (see Fig. 4 and Fig. 5). The CLANS graph is created by comparing sequences using pairwise alignments from BLAST or pairwise structure comparisons using TM-align. The CLANS sequence and structure similarity networks confirm the characteristics of ART-like superfamily of proteins, i.e., high sequence divergence (Fig. 4A) – even within a single clade – and a highly conserved tertiary structure core (Fig. 4B), also between clades. Even well-characterised ART proteins from the ECOD40 database do not always show sequence similarity high enough to be included in the graph, even with a very relaxed threshold: E-value = 1 (Fig. 4A). It is also apparent that there is more similarity within clades than between them, both when comparing sequences and structures. The small HHh clade, the minimal version of the ART-like fold (only six strands, without any extended inserts), locates centrally in the structural CLANS graph. Our analysis showed that the novel ART-like families: 1) form separate clusters of proteins and 2) generally do not group together with established, well-studied ARTs, thus confirming their distinct family status within the ART-like superfamily.

**Figure 5.**
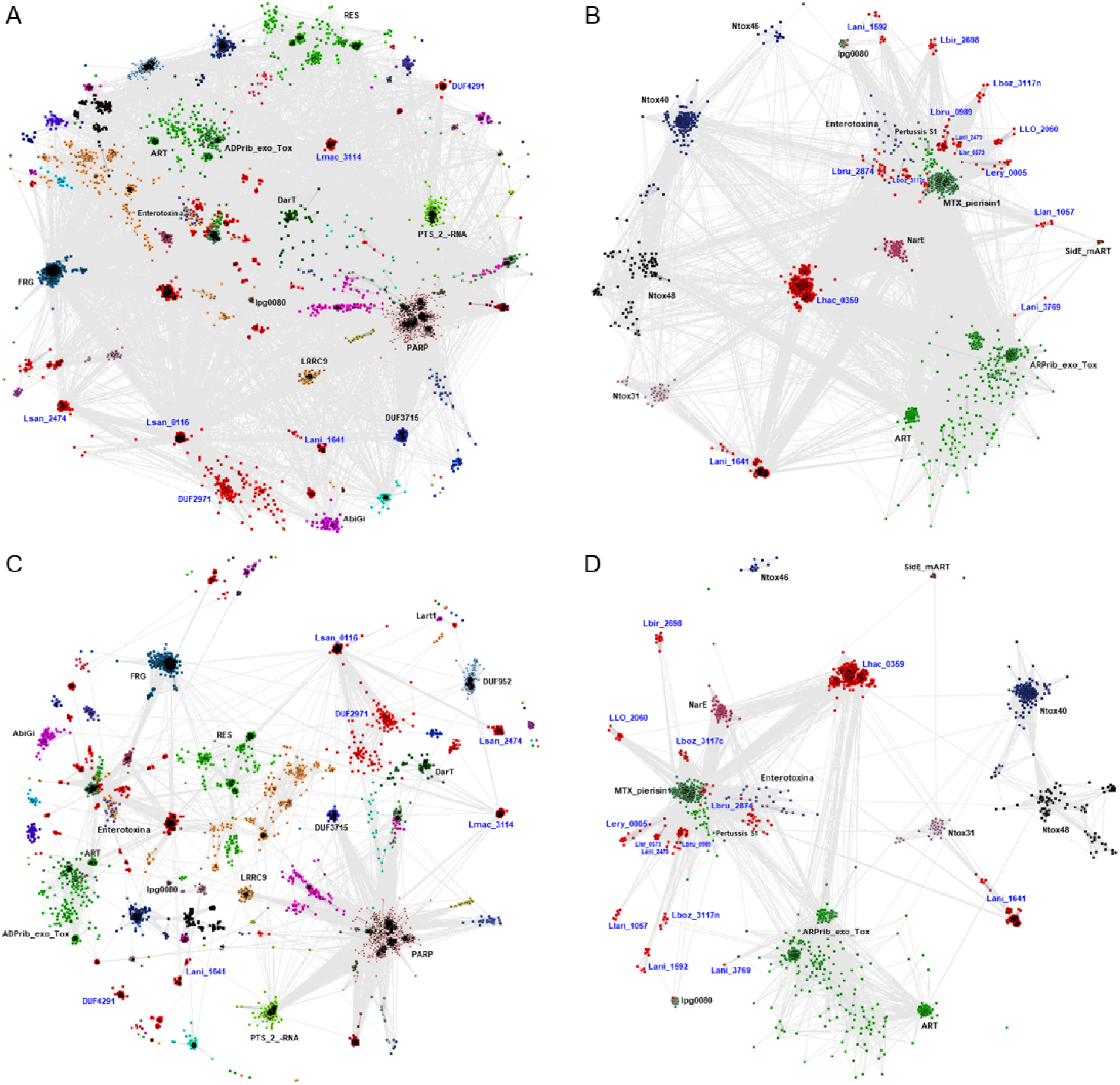
Sequence similarity relationships between new and known ART families. A and C: all ARTs families; B and D: subset of families that are closely similar to *E. coli* heat-labile enterotoxin (LT). The CLANS graph shows the sequence similarities obtained using pairwise BLAST comparisons, taking into account significant and borderline significance similarities: up to the E-value of 1 (A and B) and up to the E-value of 0.01 (C and D). Novel families of ARTs are marked in red. Colouring by families. Labelling by Pfam identifiers of selected known ART families and the gene symbols representing novel families not yet described in the Pfam database.

In-depth sequence similarity analyses (Fig. 5) compare all novel ART families (including those not described in detail in this article) with known ART-like families. The separation of the novel families (indicated in red) is clearly visible. Some of the ARTs show similarity to *E. coli* heat-labile enterotoxin (LT) (Fig. 5B and Fig. 5D).

Out of the twenty-six novel *Legionella* ART-like families, we characterised in more detail the six most interesting ones, considering breadth of their taxonomic distribution, numbers of representatives, features of operons and the conservation of key motifs. The results from this study are compiled into an ART database available at http://bioinfo.sggw.edu.pl/astarte/. This resource combines the new ART families with already known ones. In total, we present data on 82 ART families. We provide sequences, HMMs, sequence logos, 3D structure models, and brief descriptions.

### The DUF2971 family

The putative effector protein lpg1268 from *Legionella pneumophila* subsp. *pneumophila* and homologs from 13 other *Legionella* species are predicted as a novel ADP-ribosyltransferase family. The DUF2791 domain (DUF, domain of unknown function) shows statistically significant sequence similarity to known ARTs, especially AbiGi, DarT, and Tox-ART-HYD1 families (see Fig. 2, Fig. 4 and Fig. 5). DarT is the toxin element of a toxin-antitoxin (TA) system. It is an enzyme that specifically ADP-ribosylates thymidine nucleotides on ssDNA in a sequence-specific manner (Jankevicius et al., 2016; Schuller et al., 2021).

The ADP-ribosyltransferase catalytic core appears to be well conserved in this family, although the histidine residue in the catalytic motif I is poorly conserved. There is a strictly conserved tyrosine residue in motif II and two conserved glutamic acid residues in the ExE motif III.

The DUF2971 family is present in 5981 species (UniProt database), mainly in Bacteria (5913 species), Archaea (56 species), Eukaryota (6 hits, mostly in in fungi) and in *Caudoviricetes* - tailed bacteriophages (12). Among the DUF2971-possessing bacteria, one numerous group consists of soil-living saprophytic *Clostidriales* that ferment plant polysaccharides (82 species). Another large group are ubiquitous bacteria of the order *Enterobacteriales*, abundant in the human large intestine, also on the skin and in the oropharynx, or free-living in water. Most are opportunistic pathogens that infect the organisms with the weakened immune system, like *Salmonella*, *Escherichia coli*, *Klebsiella* or *Shigella*. Notably, DUF2971 homologues can be found in strains pathogenic to humans in *Escherichia coli* O157: H7 (EHEC), *Salmonella enterica*, *Vibrio cholerae*.

The appearance of the active site and the structure of the protein (the core is built of 6 beta sheets with inserts between) leaves no doubt this family should be included in the HYE clade (see Fig. 2, Suppl. Fig. 1 and Suppl. Fig. 2). The structural divergence from the HYE clade seen in the CLANS analysis (see Fig. 4) is due to unusually large helical inserts between β-strands 1 and 2, as well as between β-strands 4 and 5, that do not disrupt the core of the structure (see Suppl. Fig. 1).

About 18% of proteins having a DUF2971 domain may function as bacterial effectors (Fig. 8 and Suppl. Table S7). Predicted DUF2971 effectors are found in well-known pathogenic strains of diverse bacteria: *S. enterica*, *V. parahaemolyticus*, *V. cholerae*, *L. pneumophila* (Suppl. Table S7), suggesting that DUF2971 may play a role in their pathogenic mechanisms.

The DUF2971 domain proteins are, in a few instances, found in operons together with peptidase C26 and low affinity iron permeases. We propose that these proteins may interact in response to oxidative stress, regulation of iron metabolism and protein degradation. Under stress, such as iron excess, peptidase C26 can degrade poly-gamma-glutamate, which binds iron ions, reducing their toxicity. The degradation of poly-gamma-glutamates provides glutamic acid, a precursor to important metabolites, including glutathione, which protects against oxidative stress. Peptidase C26 can influence the availability of metabolites and iron ions by modulating the activity of ADP-ribosyltransferases in response to stress or cell damage. Iron permease regulates the transport of iron ions into the cell, and degradation of poly-gamma-glutamate by peptidase C26 affects intracellular iron levels, which is crucial for homeostasis and protection against oxidative stress. The operon-coordinated response to oxidative stress and regulation of iron metabolism promotes bacterial adaptation to changing environmental conditions.

In another interesting operon variant, we found DUF2971 together with Abi_C, KfrA_N, and a dermonecrotic toxin of the papain-like fold (see Fig. 7). The grouping of these genes in a single operon suggests a coordinated response to various forms of stress, such as bacteriophage infections, competitive pressure, and the regulation of plasmid replication.

ADP-ribosyltransferase (ART) may be involved in stress response and DNA repair. The Abi_C domain induces abortive infection, which leads to the death of the infected cell and prevention of bacteriophage spread. KfrA_N suggests involvement in the control of plasmid replication, which is crucial for the propagation of resistance genes. Papain-like fold toxins can destroy competing bacteria or protect against pathogens, which is vital in competitive environments. The integration of DNA repair mechanisms, bacteriophage defence, plasmid control, and toxin production indicate a complex system of defence and adaptation, enhancing the bacteria’s chances of survival in changing conditions.

### Lsan_0116 – an effector family common to bacteria and archaea

The Lsan_0116 novel ART-like family, named after the cognate *Legionella santicrucis* protein, shows a noticeable similarity (13% sequences identity by HHpred) to the previously described DUF2971 (see Fig. 4, CLANS diagram). However, the two families present different types of active sites: Lsan_0116 presents an RSE motif, while DUF2971 presents a modified HYE motif (see Fig. 2). Therefore, we consider them to be separate families, even though distant homologs of the two families overlap. We discovered it mainly in bacteria with it being present in most major bacterial taxa (see Fig. 3), and also sparsely in archaeons.

The active site logo (Fig. 2) shows an arginine or lysine residue in motif I and two glutamic acid residues from motif III. Motif II, which is responsible for the groove for binding the nicotinamide ring and ribose and catalysis (Bell & Eisenberg, 1996), is the most unusual. In this family, motif II has the form of [SC]WH instead of the canonical STS. The protein structure presented by the AlphaFold model is also unusual (see Suppl. Fig. 1 and Suppl. Fig. 2) – the active site is formed by the first, third and seventh beta sheets (1-3-7 instead of the typical 1-2-5 layout). The structural divergence from the RSE clade is evident in the CLANS analysis (see Fig. 4) and is due to unusual sequence insertions within the catalytic core sequence.

Majority of the Lsan_0116 family (80%) are predicted effectors, in contrast to the related DUF2971 family which has only 18% likely effectors (Fig. 8 and Suppl. Table S7).

**Figure 6.**
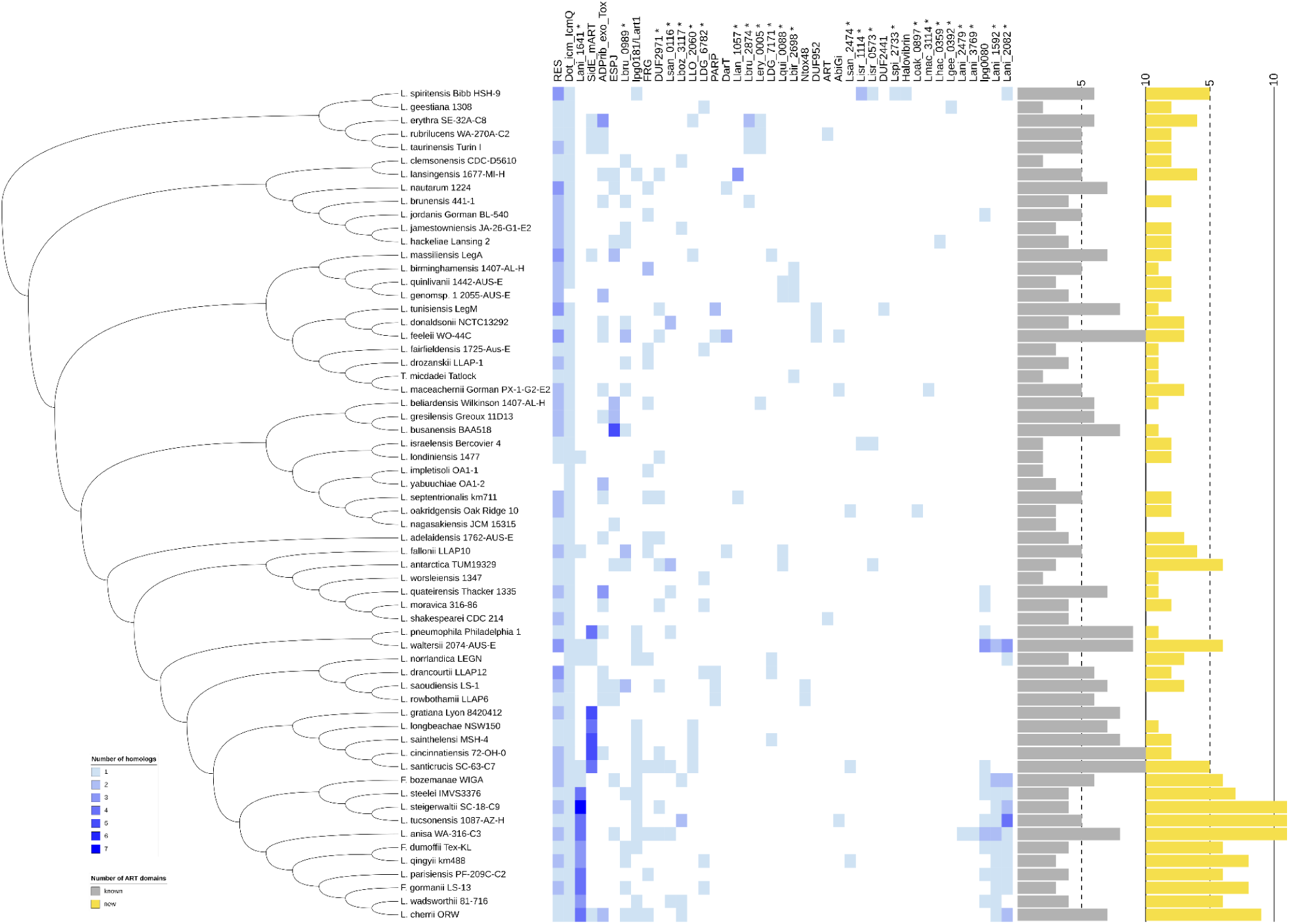
Distribution of ART families in *Legionella* strains. Numbers of homologues of novel and known ADP-ribosyltransferase families in selected members of the family *Legionellaceae*. Histograms on the right: numbers of members of known (grey) and novel (yellow) ART families identified in each species. New families are marked with asterisks above the heatmap. The DUF4291 family was not included in the diagram because *Legionella* homologs were not found in the RefSeq database (see Methods).

### The DUF4291 family – the largest atypical novel ART family, present also outside *Bacteria*

In addition to sequence searches for novel ARTs, we performed a structural search among RoseTTAFold models of all families in the Pfam database (see Methods). A single domain of unknown function (DUF4291), present in *Legionella*, stood out as significantly similar by structure to known ARTs. Based on structural similarity, conserved sequence motifs in the beta sheets 1, 2 and 5, an ART-like active site could be proposed (Fig. 2).

According to our survey, the DUF4291 is a novel effector family with similarity to ARTs. According to the Pfam database annotations, there are 2 conserved motifs in this uncharacterized family. Structural alignment suggests that these motifs are atypical counterparts of the motifs I and II of the ART catalytic site (QAY and WVK correspond to HxT and STS, respectively).

Although the predicted active site in this family has a very unusual appearance, the catalytic motif III includes a “classical” glutamine. The new family is strikingly well conserved. The DUF4291 domain shows statistically significant structure similarity to HYE clade of ARTs, especially Pfam families Exotox-A_cataly and RES. We could not identify significant sequence similarity to any known family of ARTs. (Suppl. Fig. 1 and Suppl. Fig. 2). Also, the correct docking of NAD to the obtained model suggests that the DUF4291 domain may have ART activity (Suppl. Fig. 2).

The DUF4291 family is present in 4540 species, mainly in *Bacteria* (3887 organisms) and in *Eukaryota* (646 species). This family is observed in *Legionellaceae* (it has only been identified in the *Tatlockia* genus which is closely related to the genus *Legionella* (Tindall, 2020; Saini & Gupta, 2021)) and in other, well-studied bacteria e.g., *Escherichia coli* O103:H25, *Salmonella enterica*, *Xanthomonas oryzae*, *Myxococcus xanthus*, *Klebsiella aerogenes*. In bacteria, most hits were identified in bacterial phytopathogens and animal pathogens, including human ones or in opportunistic pathogens, and in harmless commensals of mammals and bacteria that occur in water or soil. We also found it common in bacteria widely used to produce antibiotics and other therapeutics (e.g. *Streptomycetaceae*), in photosynthesising *Cyanobacteria* and in the human microbiome (e.g. *Firmicutes*). DUF4291 was also found in abundance in organic compound-degrading bacteria of the order *Myxococcales* and in plant-symbiotic nitrogen-fixing rhizobia.

In *Eukaryotes,* this family is present mainly in *Fungi* (605 hits), particularly *Dikarya*. It is widespread in *Aspergillaceae.* It also occurs in saprophytic fungi of the genus *Fusarium*. In *Metazoa,* the DUF4291 family was found in some taxa *Animalia,* among others in segmented annelid worms, urchins, lancelets and molluscs, but is absent from arthropods, tunicates and vertebrates. In addition, DUF4291 is found in *Alveolata*, *Amoebozoa*, *Rhodymeniophycidae* and *Viridiplantae*. Proteins from the new family are occasionally found in *Archaea* and *Viruses*.

Genomic neighbourhood analysis of DUF4291 family proteins (Fig. 7) revealed the common presence of TetR_N and NUDIX domains (in 28% and 18% of analysed neighbourhoods, respectively). TetR_N is a domain found in several bacterial and archaeal transcriptional regulators. The NUDIX hydrolase domains are widespread and usually function as pyrophosphohydrolases. Some NUDIX proteins degrade potentially mutagenic, oxidised nucleotides while others control the levels of metabolic intermediates and signalling compounds. Slightly less frequent are Macro and ADP_ribosyl_GH domains, co-occurring in 9-10% of DUF4291 neighbourhoods. Macro and ADP-ribosylglycohydrolase (ADP_ribosyl_GH) domains are elements of ADP-ribosylation signalling and complete the “writer-reader-eraser” triad. Macros can act as “readers’’ or “erasers”, whereas ADP-ribosylhydrolyses are “erasers” of ADP-ribosylation marks. Another known ART domain – RNA 2’-phosphotransferase (PTS_2-RNA) was observed almost as often (10%) in the genomic neighbourhoods of DUF4291. Taken together, these observations strongly support involvement of DUF4971 in ADP-ribosylation signalling and transcription and/or RNA biochemistry.

**Figure 7.**
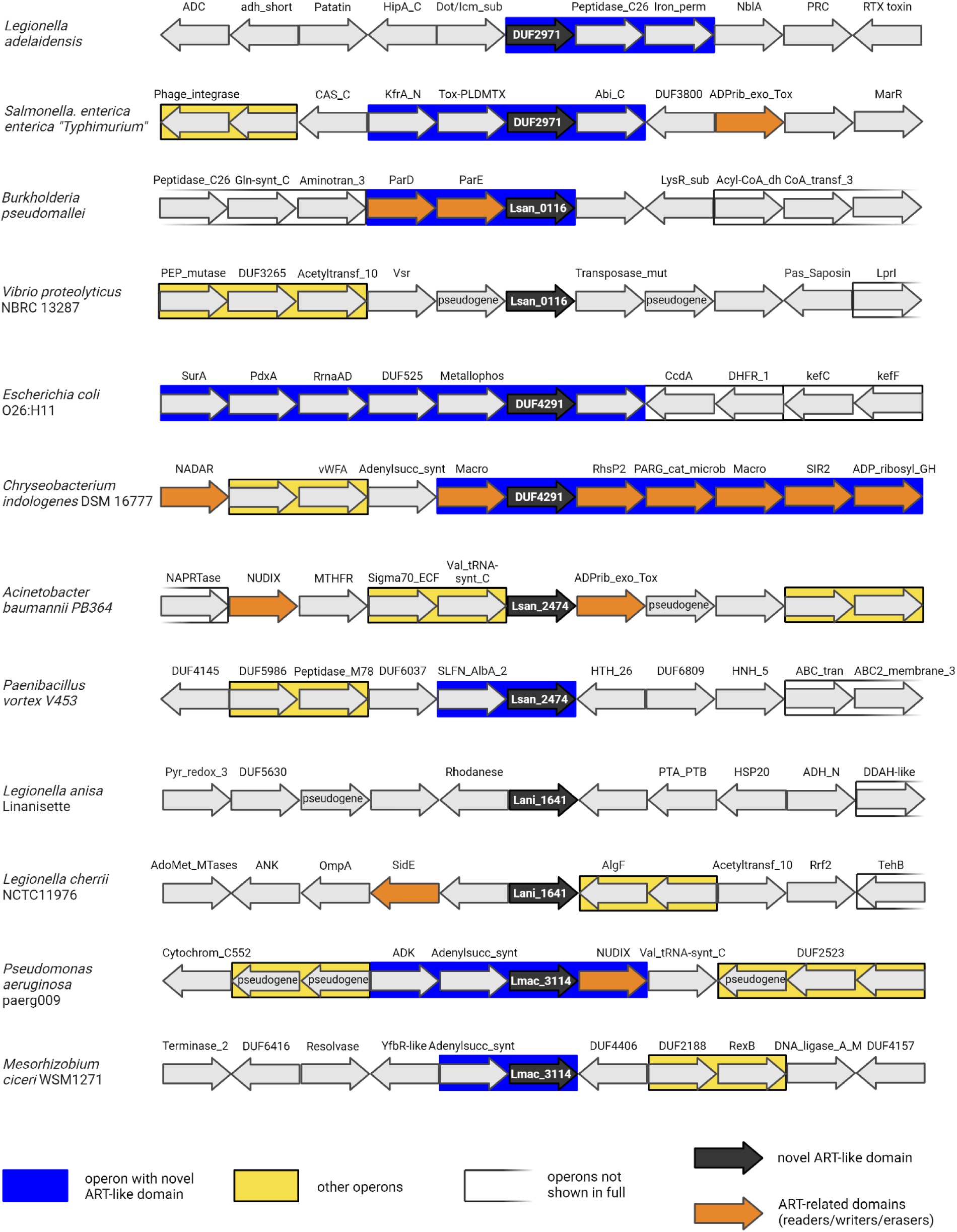
Genomic neighbourhoods of selected novel ART-like domains. Pfam/InterPro protein domain short names or gene names used. The known or predicted operon information was obtained from the BioCyc database. Created with BioRender.com

It is estimated that more than 40% of proteins having a DUF4291 domain may be effectors (see Fig. 8 and Suppl. Table S7). However, due to the prediction algorithm using the presence of eukaryotic-like domains (ELDs), there is a risk of overprediction.

### Lsan_2474, a new ART-like domain, similar to AbiGi

This novel family, named after the *Legionella santicrucis* Lsan_2474 protein, is a distant homologue R-S-E clade families, AbiGi (8% sequence identity) and DUF2971. FFAS sequence alignments identified three conserved active site motifs typical of the R-S-E clade (R-SxS-ExE), however, motif II is often mutated to CxS. The presence of the novel family was confirmed only in Bacteria (133 organisms) from diverse taxa: *Gammaproteobacteria, Betaproteobacteria, Epsilonproteobacteria*, *Terrabacteria* group, *Acidobacteria*, and *Bacteroidetes*, e.g., *Vibrio cholerae, Pseudomonas syringae, Yersinia enterocolitica* and a virus (Siphoviridae). There are bacterial effectors in the Lsan_2474 family and we estimate that about 78% of proteins having a Lsan_2474 domain may be effectors (Fig. 8 and suppl. S7).

### Lani_1641, a new ART-like domain, similar to Ntox31 and Enterotoxin_a

Lani_1641 is a small, novel ART-like family named after *L. anisa* protein Lani_1641. Although the active site presents the RSE motif, sequence comparisons (Fig. 4 and Fig. 5) do not place this family close to the RSE clade families. The matching of the protein structure model to known ART structures shows strong similarity to Pertussis toxin, Scabin and PARPs with 11-18% sequence identity and significant DALI Z-scores between 5.5 and 11.1. The CLANS analysis also places this family in the RSE clade, and the protein model suggests the presence of a catalytic core composed of seven beta sheets.

Members of this family were detected only in *Legionellaceae*. It is estimated that about 65% of proteins having a Lani_1641 domain may be effectors (Fig. 8 and Suppl. Table S7). The ART-like Lani_1641 domains are not accompanied by other functional and structural domains in proteins. Also, no multigene operons involving the Lani-1641 domain were identified (Fig. 7).

### Lmac_3114 new ART-like domain, similar to DarT

The novel Lmac_3114 family, named after *Legionella maceachernii* protein Lmac_3114 is a distant homologue of DarT domain – from the H-Y-E clade of ART. In Lmac_3114, the catalytic motif II [Y-x-x] contains a conserved tryptophan residue. DarT family members act as the toxins in toxin-antitoxin (TA) system, by specifically modifying thymidine nucleotides on ssDNA.

Its closest relatives identified by FFAS, HHpred and Phyre2 also include the ART family RNA 2’-phosphotransferase. Sequence alignments identified three conserved active site motifs (Hxx, FFW, and xxE). Structural comparisons do not make it possible to determine precisely to which clade this new family should be assigned. Greatest similarity is found to the DarT family from the HYE clade and PTS_2-RNA from the HHh clade, however, the presence of a conserved glutamic acid residue in the third motif leads us to assign it to the HYE clade. For both clades, the presence of six beta-sheets in the catalytic core is characteristic, which is reflected in the Lmac_3114 model.

This family is detected only in bacteria from *Gammaproteobacteria*, *Alphaproteobacteria*, *Betaproteobacteria*, *Deltaproteobacteria*, *Nitrospirae*, *Terrabacteria* group, *Acidobacteria*, PVC group and *Bacteroidetes*, e.g., *Pseudomonas aeruginosa*, *P. syringae, Xanthomonas oryzae* and *Burkholderia pseudomallei*.

Genomic neighbourhood analysis of Lmac_3114 family proteins provided evidence that the adenylosuccinate synthetase domain is extremely common in its vicinity (co-occurrence in 60% of cases) and, in at least some cases, forms an operon with the ART-like domain (Fig. 7). The presence of genes encoding ADP-ribosyltransferase and adenylosuccinate synthase in the operon suggests their functional association, e.g. co-operation in specific metabolic or regulatory processes. We have identified such an operon in nitrogen-fixing soil bacteria (*Rhizobium azooxidifex* and *Mesorhizobium ciceri*). A more elaborate version of the operon is found in *Pseudomonas aeruginosa* – there the ART-like domain is found together with the AAA domain, adenylosuccinate synthetase and NUDIX domain, a hydrolase that cleaves nucleoside diphosphates linked to another moiety. The presence in the common operon of adenylosuccinate synthase and ART domains suggests regulation of ART activity by changes in purine synthesis, e.g. indirect modulation of ADP-ribosyltransferase activity in either cellular regulation or stress response by regulating NAD+ availability.

### Analysis of potential NAD binding sites

To evaluate the compatibility of the novel ART families with NAD binding, we used HADDOCK to dock NAD to reference protein structure models for families discussed in detail in this article. For comparison, we performed re-docking for a crystallographic complex consisting of NAD bound to eukaryotic mono-ADP-ribosyltransferase ART2.2 (PDB code 1OG3) (Ritter et al., 2003). Among the six novel ART families, the best NAD docking scores ranged from −60.2 to −31.5 while the score for redocking to the NAD bound crystal structure, the score was −49.6. The best scores (under −60.0) were obtained for Lsan2474 and DUF4291 representatives. In all cases, manual identification of amino acids forming the putative NAD pocket based on sequence conservation and structural alignments to known ARTs resulted in a more favourable docking than automatic detection of ligand-binding residues. In almost all cases (except Lmac_3114), the docking procedure resulted in the NAD ligand in a correct position, i.e. resembling the NAD poses observed in known ART crystal structures (Suppl. Fig. 1). Intriguingly, for Lmac_3114, the NAD molecule is located in a non-standard location, however, the predicted active site residues interact with the bound NAD molecule which may suggest an atypical catalytic mechanism, different from most ART families. Mapping sequence conservation onto the protein surfaces shows that highly conserved residues group within and near in the predicted NAD binding pockets. Close-up view of the NAD sites focusing on conserved residue side chains shows that the poses of NAD molecule are very similar for all models, with adenine ring stacking out (again except of Lmac_3114).

The active site signatures indicated by sequence logos (Fig. 2) were confirmed by structural models and NAD docking results as likely catalytically relevant residues involved in binding the NAD molecule (Suppl. Fig. 1, Suppl. Fig. 2 and Suppl. Table S8). We obtained similar results using the AlphaFold3 server (Abramson et al., 2024) to model representative proteins from each family with NAD+ (data not shown). These observations support our hypothesis that the newly discovered ART-like families can bind NAD and be functional ADP-ribosyltransferases.

To gain insight into potential functions of the novel ART-like families, we predicted the content of secreted proteins and effectors in each family by analysing presence of signal peptides, the amino acid composition of the C-terminus of the sequence, the binding site of the secretion chaperone protein and the presence of eukaryotic-like domains (see Methods and Suppl. Table S7). Among the families discussed in detail in this article, all are predicted to include secreted effector proteins. Particularly prominent is the Lsan_0116 family, in which we predict that more than 80% of the proteins will be effectors (Fig. 8).

**Figure 8.**
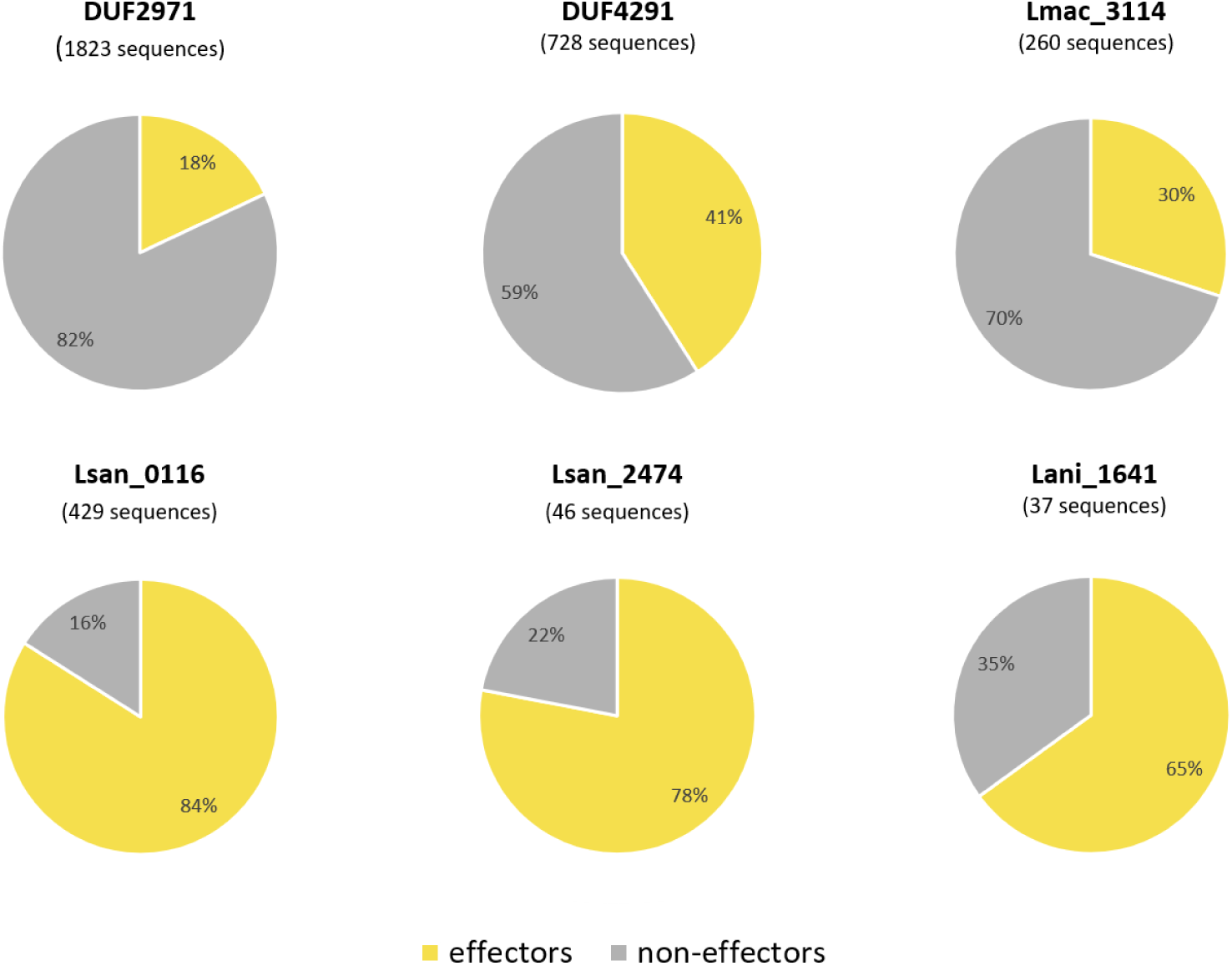
Predicted share of effector proteins in the novel ART families. Predictions made using SignalP 6.0, EffectiveDB and BastionX. Data for all identified families available in the supplement (Suppl. Table S7).

## Discussion

In our bioinformatic examination of 41 *Legionella* species, we have documented 63 groups of *Legionella* orthologues with convincing sequence or structure similarity to ADP-ribosyltransferases, that can be organised into 39 ART-like families. Among these families, 26 families are novel ART superfamily members (see Table 1). One of the novel families (DUF2971) is represented in *L. pneumophila.* The newly identified ART families, initially found by sequence searches, have been further corroborated by artificial intelligence-driven structural predictions.

**Table 1.**
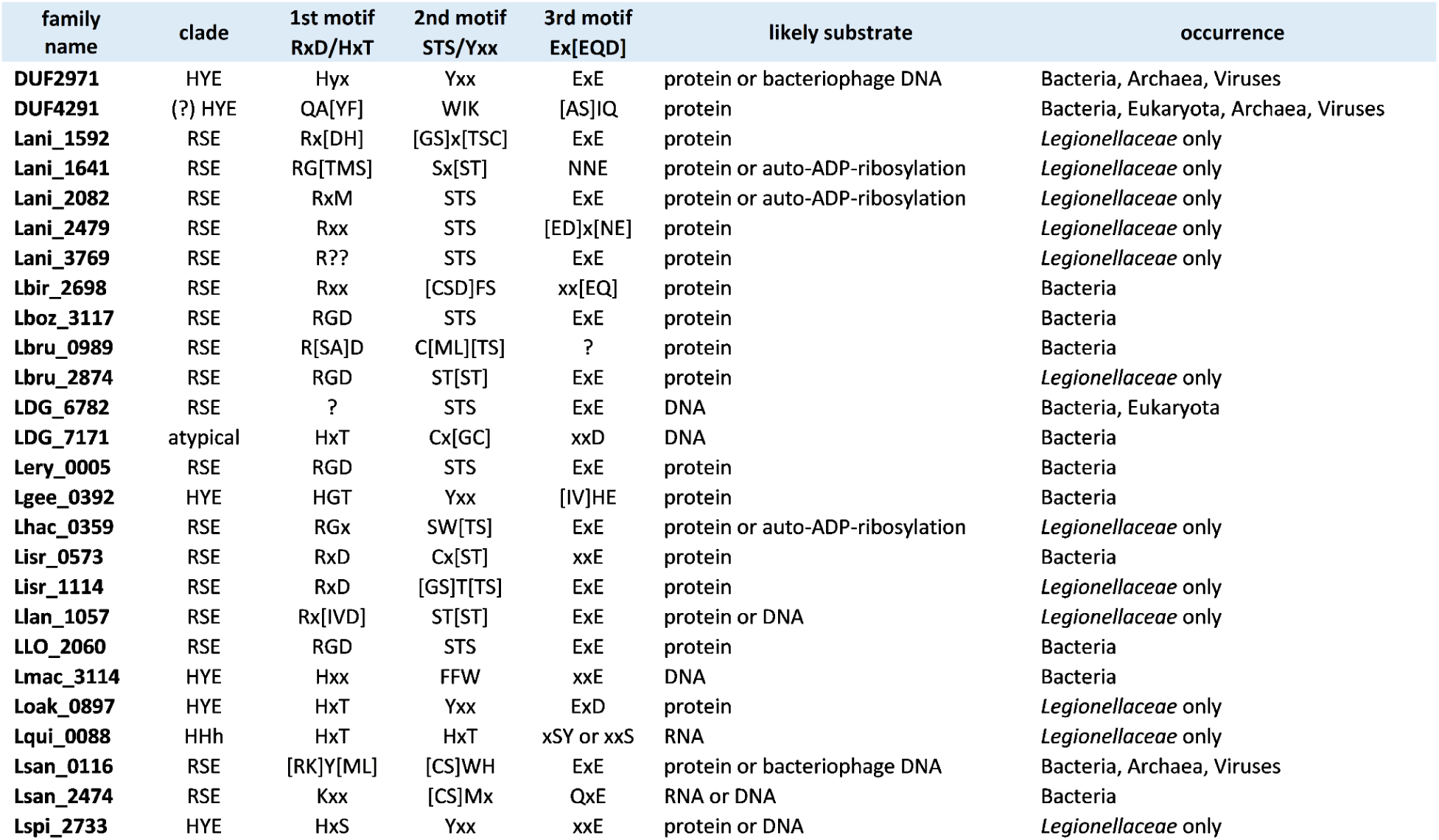
Novel ADP-ribosyltransferase families identified in the *Legionella* pan-proteome. Families are identified by Pfam names (DUF) or by symbols of representative genes (Burstein et al., 2016).

The DUF2971 family shows similarity to DarT proteins and scabin, a DNA-acting ADP-ribosyltransferases (Lyons et al., 2016) suggests that DUF2971 effectors could modify host or invader’s DNA.

The largest newly identified ART-like family, DUF4291, including approximately 5 000 members, shows structural and sequence similarity to the diphtheria toxin family of mono-ART toxins, e.g. *P. aeruginosa* exotoxin A, *C. diphtheriae* diphtheria toxin and cholix toxin from *V. cholerae* (Jørgensen et al., 2008). These ARTs are important virulence factors belonging to the class of exotoxins secreted by pathogenic bacteria and cause diphtheria, cholera and pneumonia, respectively. All of them ADP-ribosylate elongation factor 2 in host cells, so it is plausible to hypothesise that the DUF4291 family modifies proteins rather than nucleic acids.

The ART complement of the *Legionella* genus showcases diverse effector ARTs, possibly corresponding to adaptability to various hosts. For example, the extensively studied *Legionella pneumophila*, possesses genes known to facilitate infection in various hosts, which exhibit both similarities and distant relationships with eukaryotic ARTs, possibly acquired through horizontal gene transfer.

Although the predicted effector functions remain to be validated, these novel ART-like families present attractive candidates for experimental studies into mechanisms of pathogenicity due to their likely crucial roles in the pathogen’s survival. While this manuscript was being finalised, a thorough study presented a detailed survey of structural domains in all the effectors of *L. pneumophila* (Patel et al., 2024). While our study addresses only ART-like proteins, it is complementary to work of Patel et al. because we surveyed 41 species instead of one and did not limit the exploration to effectors.

In summary, our survey of the *Legionella* ADP-ribosyltransferase world provides insights into the pathogen’s toolkit for infection and offers a broader perspective on nature’s remarkable adaptability and ingenuity in the use of the evolutionarily successful ART-like superfamily. The results from this study are available at http://bioinfo.sggw.edu.pl/astarte/.

## Acknowledgements

We thank Drs A. Muszewska, V. Tagliabracci and B. Mayro for critical reading of the manuscript and helpful comments.

## Author contributions

**MK**: conceptualised the project, performed the experiments, analysed the data, prepared figures and tables, authored or reviewed drafts of the paper, and approved the final draft

**BB**: performed the experiments, analysed the data, and approved the final draft.

**MG**: performed the experiments, analysed the data, prepared supplemental tables, and approved the final draft.

**KP**: conceptualised the project, co-supervised the project, authored or reviewed drafts of the paper, and approved the final draft.

**MD**: conceptualised the project, co-supervised the project, performed the experiments, analysed the data, prepared supplemental figures and/or tables, authored or reviewed drafts of the paper, and approved the final draft.

## Funding and additional information

K.P. was supported by the Polish National Science Centre grant 2019/33/B/NZ2/01409. M. G. was supported by the Polish National Science Centre grant 2019/35/N/NZ2/02844. The funders had no role in study design, data collection and analysis, decision to publish, or preparation of the manuscript.

